# A conserved, immune-regulated peritrophin promotes *V. cholerae* colonization of the arthropod intestine

**DOI:** 10.1101/2025.04.21.649765

**Authors:** Daniela Barraza, Tania F. Paulo, Lauren Findley, Saiyu Hang, Paula I. Watnick

## Abstract

*Vibrio cholerae* is a human diarrheal pathogen and an estuarine organism that associates with both terrestrial and aquatic arthropods. Using the model terrestrial arthropod *Drosophila melanogaster*, we previously showed that *V. cholerae* forms a multi-layered bacterial structure called a biofilm in the arthropod intestine and activates the arthropod intestinal innate immune response. Here we show that activation of the immune response in enterocytes decreases *V. cholerae* colonization of the arthropod intestine, while activation of the immune response in enteroendocrine cells that express the enteroendocrine peptide tachykinin (Tk) promotes *V. cholerae* colonization. To uncover the basis of this observation, we measured the impact of *Tk*^RNAi^ on intestinal gene expression by RNA-seq analysis. In addition to increasing expression of antimicrobial peptides and lipases, Tk activated the expression of chitinases and chitin-binding proteins. These proteins interact with chitin fibrils in the peritrophic matrix (PM), a protective coating that overlies the arthropod intestinal epithelium. One of these Tk-activated PM components, the small, secreted chitin-binding protein Peritrophin 15a (Peri-15a), is essential for robust *V. cholerae* colonization of the gut. Homologs of Peri-15a are widespread in both terrestrial and aquatic organisms including marine non-biting midges, marine copepods, rotifers, and cyanobacteria. We propose that Peri-15a and its homologs, found in the intestines of diverse arthropods, either serves as a receptor or reveals a PM epitope that promotes *V. cholerae* attachment to the intestinal surface. Therefore, activation of the enteroendocrine cell intestinal innate immune response by *V. cholerae* may, in fact, represent a colonization strategy.

## Introduction

*Vibrio cholerae* is best known as a human pathogen that causes the diarrheal disease cholera due to expression of the principal virulence factors, the toxin co-regulated pilus and cholera toxin (1). However, both toxigenic and non-toxigenic strains of *V. cholerae* are found in estuarine aquatic environments (2–5). In these environments, *V. cholerae* is found in association with large and small marine arthropods including midges, rotifers, copepods, and cladocerans (4–7). The exoskeletons of these organisms are rich in chitin, and some have suggested that the ability of *V. cholerae* to attach to, degrade, and grow on chitin contributes to this environmental association (8–12). Chitin also activates natural competence in *V. cholerae*, thus promoting horizontal transfer of genetic material, which is likely important to environmental adaptation and survival (13, 14).

*Drosophila* melanogaster, the common fruit fly and model arthropod, has revealed novel aspects of the interaction of *V. cholerae* with both the arthropod and mammalian intestinal epithelia (15–23). The *Drosophila* intestine consists of a foregut, midgut, hindgut and rectum (Fig 1A). The midgut is the principal site of digestion and absorption of nutrients, much like the mammalian small intestine and is separated from the intestinal lumen by a porous peritrophic matrix (PM) (24). The PM consists of a lumen-exposed web-like structure comprised of chitin and chitin-binding proteins known as peritrophins and an inner layer of mucins that are anchored to the chitin (25). The midgut is divided into the cardia and anterior midgut (AMG) where the microbiota reside, the middle midgut that is acidic, and the posterior midgut (PMG), which is relatively microbe-free. Because microbes entering the midgut first encounter the AMG, the innate immune response in this region is critical for intestinal defense. Cell types in the midgut include enterocytes (ECs) with interspersed intestinal stem cells and enteroendocrine cells (EECs). ECs absorb nutrients from the diet, while stem cells replenish the intestinal epithelium as needed and EECs respond to soluble signals from the lumen and host by releasing packets of small bioactive peptides known as enteroendocrine peptides. EECs in the AMG are specialized for responding to microbial metabolites (26). This, in addition to microbe-associated molecular patterns (MAMPs) such as peptidoglycan, may signal the presence of viable microbes in the gut (27, 28).

**Figure 1:**
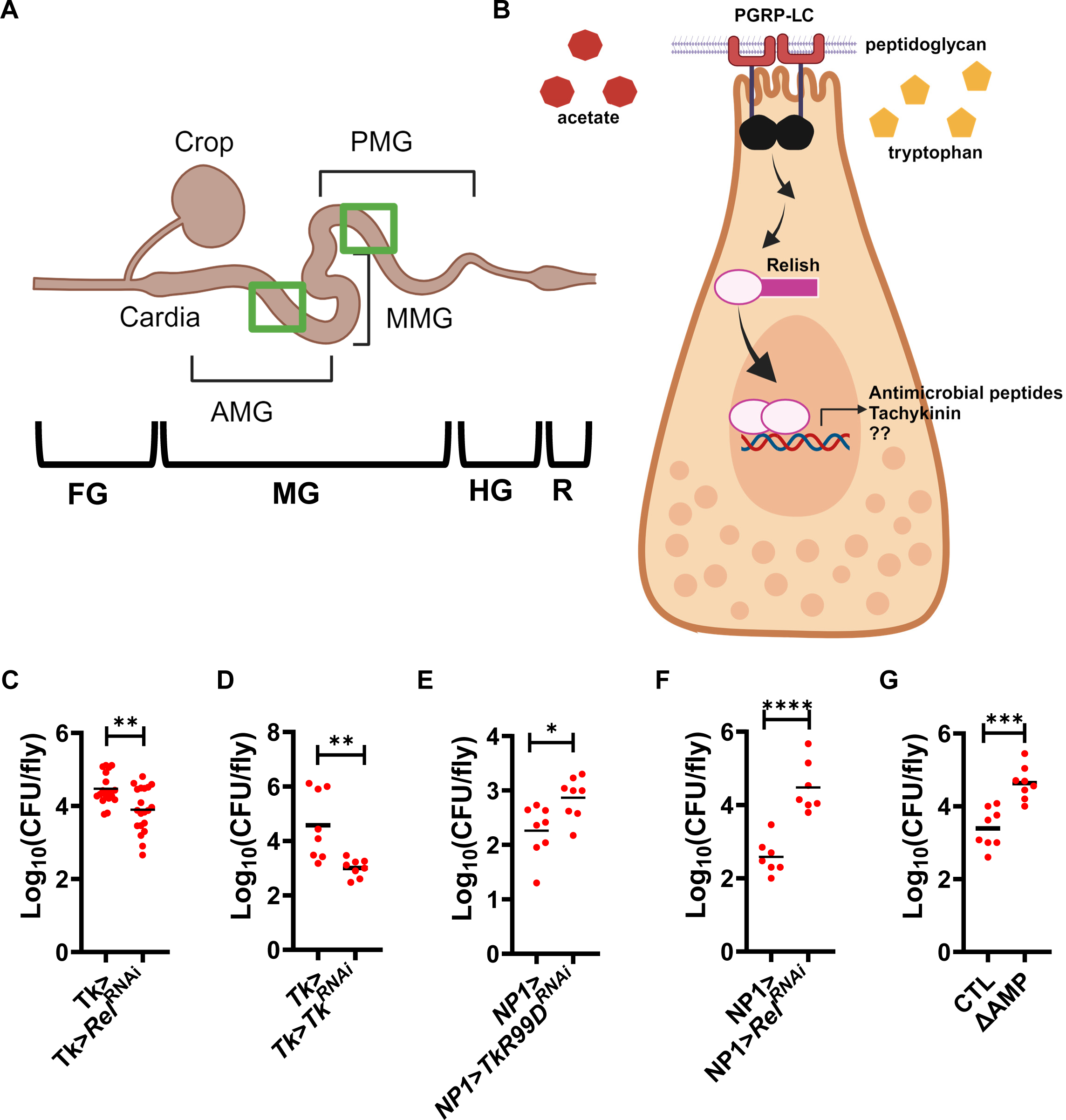
Knockdown of IMD and Tk signaling in EECs decreases *V. cholerae* colonization of the arthropod intestine, while knockdown in ECs increases colonization. (A) Schematic of the different regions of the *Drosophila* intestine including the foregut (FG), midgut (MG), hindgut (HG), and rectum (R). The midgut is comprised of the anterior midgut (AMG), the middle midgut (MMG), and the posterior midgut (PMG). Green boxes indicate the regions of the AMG and PMG where lipid content is assessed. (B) IMD pathway activation signals and signaling in Tk-expressing enteroendocrine cells of the AMG. Quantification of *V. cholerae* intestinal colonization of (C) Tk+ EEC driver control (Tk>) and Tk>*Rel*^RNAi^ flies, (D) Tk+ EEC driver control (Tk>) and Tk>*Tk*^RNAi^ flies, (E) NP1> and NP1>*TkR99D*^RNAi^ flies, (F) NP1> and NP1>*Rel*^RNAi^ flies, and (G) parental *w*^1118^ (CTL) and flies with 10 AMPs deleted (ΔAMP). Flies were fed *V. cholerae* for 48 hours, and non-adherent bacteria were then washed out by feeding on PBS for 24 hours. The mean of at least 6 flies is shown. A student’s t test was applied to log-transformed data to calculate significance. **** p<0.0001, *** p<0.001, ** p<0.01, * p<0.05.

Each cell type in the intestinal epithelium encodes components of the innate immune immunodeficiency (IMD) pathway (Fig 1B) (29). The IMD pathway responds to peptidoglycan signals by phosphorylating and cleaving the transcription factor Relish. The N-terminus of Relish then translocates to the nucleus where it activates transcription of antimicrobial peptides (AMPs) such as Diptericin A (DptA), Attacin A (AttA), and Cecropin A1 (CecA1) (30). EECs of the AMG that express the enteroendocrine peptide Tachykinin (Tk) deploy IMD signaling in response to small molecules such as acetate and tryptophan (19, 27, 28). In these EECs, IMD signaling activates transcription of *Tk*, which promotes intestinal lipid utilization in addition to AMP expression (28, 31). In the absence of an invasive intestinal infection, the response of the intestine to microbes is principally under the control of enteroendocrine cells and Tk (26).

*Drosophila* ingestion of *V. cholerae* results in the formation of a biofilm in the posterior *Drosophila* midgut without further invasion and activates IMD signaling in EECs leading to an increase in intestinal Tk and AMP expression (17, 19, 27). Here we report that, despite the susceptibility of *V. cholerae* to AMPs, innate immune signaling in EECs increases *V. cholerae* colonization of the arthropod intestine. RNAseq analysis of Tk> driver control and Tk>Tk^RNAi^ flies showed that, in addition to the known impacts of Tk signaling on genes encoding AMPs and lipid catabolism enzymes, Tk activates transcription of genes encoding chitin modification and chitin-binding proteins. We demonstrate that a small chitin-binding protein, Peritrophin-15a (Peri-15a), which consists of a secretion peptide and one type 2 chitin binding domain, is activated by *V. cholerae* infection and increases *V. cholerae* adhesion to the arthropod intestine. Underscoring the importance of our findings, the literature suggests that association of *Vibrio* species with zooplankton activates both the innate immune response and transcription of chitin-binding proteins (32). Furthermore, AlphaFold predictions show that close structural homologs of Peri-15a are present in many organisms that associate with *V. cholerae* in the environment including copepods, rotifers, cyanobacteria, non-biting midges, and house flies (33). Based on these findings, we propose that *V. cholerae* exploits the host innate immune response to increase colonization of the intestinal surface of environmental arthropod hosts.

## Results

We previously showed that *V. cholerae* infection activates intestinal IMD signaling in EECs leading to increased AMP transcription in the arthropod intestine (19, 28). We hypothesized that this response to *V. cholerae* infection would limit intestinal colonization. To test this, we fed *V. cholerae* to Tk> and Tk>*Rel*^RNAi^ flies for 48 hours and then washed out non-adherent bacteria by transferring the flies to phosphate-buffered saline (PBS) for 24 hours. Unexpectedly, *Rel*^RNAi^ in Tk+ cells decreased *V. cholerae* colonization, suggesting that IMD signaling in Tk+ EECs boosts intestinal colonization (Fig 1C). IMD signaling in EECs is essential for Tk expression in the AMG (28). To test the role of Tk in this process, we measured colonization in a Tk>*Tk*^RNAi^ fly. Again, we observed a decrease in colonization (Fig 1D). Tk communicates with enterocytes via the TkR99D receptor (27, 31). To determine whether colonization was brought about by an EC-specific product, we compared colonization of EC driver control NP1> and NP1>*TkR99D*^RNAi^ flies. *TkR99D*^RNAi^ in ECs increased *V. cholerae* colonization as did *Rel*^RNAi^ (Fig 1E and F). We reasoned that, as a principal source of AMPs, a block in EC AMP production due to either decreased Tk or IMD signaling might explain the observed increase in colonization. To test this, we compared *V. cholerae* colonization of a fly with 10 AMPs deleted (ΔAMP10, Def^SK3^, AttC^Mi^, DroAttAB^SK2^, Mtk^R1^, DptAB^SK1^; Drs^R1^, AttD^SK1^) with the parental control (Fig 1G) (34). As predicted, the ΔAMP10 fly displayed increased colonization. We conclude that AMPs diminish *V. cholerae* intestinal colonization, that a Tk-regulated product of EECs promotes *V. cholerae* intestinal colonization, and that *V. cholerae* intestinal colonization of the arthropod intestine is a composite of these two opposing processes.

To identify host colonization factors regulated by Tk, we performed RNAseq on the intestines of Tk>*Tk*^RNAi^ and Tk> driver-only control flies (Table S1 and Fig 2A). Using a padj ≤0.05 and 2-fold change as thresholds, we identified 136 differentially regulated genes of which transcription of 69 were increased and transcription of 67 were decreased. Tk is known to regulate both innate immune signaling and lipid catabolism (28, 31). Based on the FlyBase entries, RNA-seq analysis showed 20 differentially regulated genes involved in innate immunity (green circles, Fig 2A and Table S1) and 9 involved in lipid catabolism (brown circles, Fig 2A and Table S1) (35). To confirm a subset of these, we measured transcription of the genes encoding the AMP Defensin (Def) and the lipases CG17192 and Lipase 3 (Lip3). As shown in Fig 2B, these measurements corroborated the RNA-seq results.

**Figure 2:**
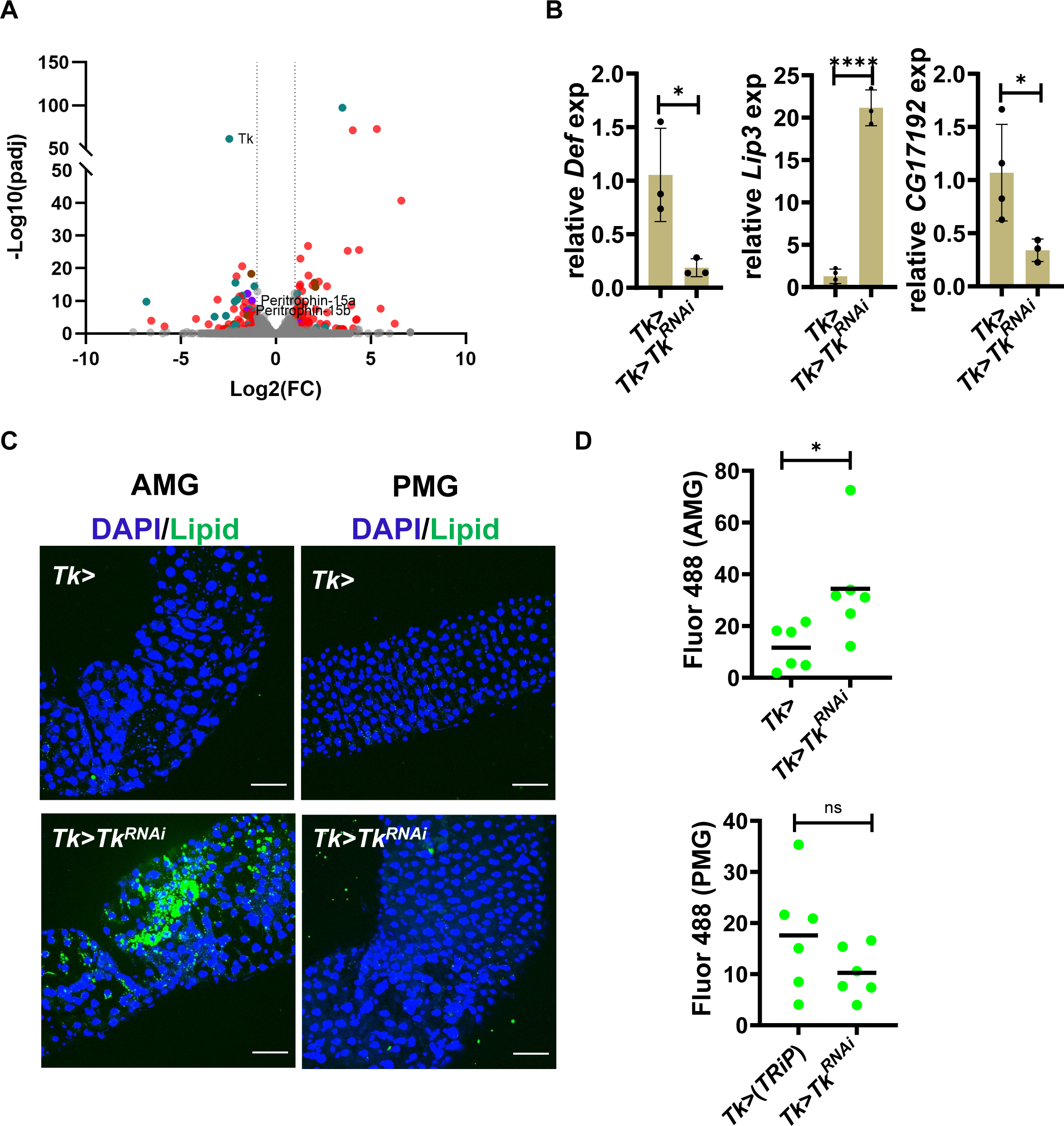
RNA-seq comparing transcription in the intestines of Tk> and Tk>*Tk*^RNAi^ flies reveals differential regulation of genes involved in the innate immune response, lipid catabolism, and chitin binding. (A) RNA-seq was performed on the intestines of Tk> and Tk>*Tk*^RNAi^ flies using biological triplicates. Volcano plot showing the log_2_fold-change (FC) and padj for gene expression in Tk>*Tk*^RNAi^/Tk> conditions. The vertical dotted lines indicate a 2-fold change. Colored points meet the threshold ≥ 2-fold change and padj < 0.05. Significantly regulated genes involved in innate immunity are shown in green (including *Defensin* or *Def*), those involved in lipid catabolism are shown in brown (including *Lip3* and *CG17192*), and those involved in chitin binding are shown in purple. (B) qRT-PCR quantification of the indicated genes in the intestines of Tk> and Tk>*Tk*^RNAi^ flies . These genes were also significantly differentially regulated in the RNA-seq experiment. The mean of biological triplicates is shown. Error bars indicate the standard deviation. A Welch’s t test was used to evaluate significance. (C) Fluorescence images showing DAPI and BODIPY (Lipid) staining in the anterior midgut (AMG) and posterior midgut (PMG) of the indicated fly genotypes. Scale bars, 50 μM. (D) Quantification of total fluorescence in the AMG and PMG of the indicated flies incubated with BODIPY. Six intestines were evaluated. The mean is shown. A students’ t-test was used to evaluate significance. **** p<0.0001, * p<0.05, ns not significant.

Opposite regulation of the two lipases *CG17192* and *Lip3* was unexpected. Therefore, we compared lipid accumulation in the intestines of Tk> control and Tk>*Tk*^RNAi^ flies. As previously reported, lipid accumulation was observed in the intestines of Tk>*Tk*^RNAi^ flies (Fig 2C and D) (31). Importantly, this was only observed in the AMG but not in the PMG. Because Tk is present in both the AMG and PMG, we conclude that control of lipid accumulation is specific to the AMG Tk (28, 31).

In addition to genes encoding metabolic, regulatory, and unknown functions, we noted four genes annotated as chitin-binding, three of which were activated by Tk and one of which was repressed. Chitin-binding proteins are structural components of the PM, the protective layer of the intestine closest to the lumen where *V. cholerae* is located during infection (36). Previous publications suggested that disruption of the peritrophic matrix increased the susceptibility of *Drosophila* to intestinal bacteria and yet we had observed a decreased burden of *V. cholerae* with *Tk*^RNAi^ (37).

Nevertheless, we hypothesized, that a chitin-binding protein might enhance pathogen colonization if it promoted adhesion to the PM surface without disrupting PM structure. A previous study of the tsetse fly showed that the peritrophins Pro 1 and 2, which are homologs of the *Drosophila* Peritrophins-15a and b (Peri-15a and b), had little impact on the structure of the PM and did not affect colonization of the intestine by the Gram-negative rods *Enterobacter* and *Serratia* (38). Because transcription of *Peri-15a* and *Peri-15b* was significantly decreased in the TK>*Tk*^RNAi^ RNA-seq analysis as compared with the Tk> control, we reasoned they might be the Tk-regulated *V. cholerae* adhesion factor we sought. Peri-15a and 15b are two related proteins consisting of 92 aa and 93 aa, respectively. They are encoded in the second chromosome near each other. An alignment using Clustal Omega showed that they are 45% identical and 58% similar (Fig 3A) (39). AlphaFold predicts very similar structures for the two proteins, which consist of an N-terminal signal peptide and one type 2 chitin binding domain (Fig 3B) (33, 40). The *Peri-15a* transcript is most highly expressed in enteroendocrine cells of the AMG, and is approximately 40X and 750X more highly expressed, respectively, than *Peri-15b* and *Peritrophin A*, the other two named peritrophins in the *Drosophila* genome (Fig 3C) (29). Therefore, we focused our studies on Peri-15a. We first confirmed our RNAseq results in an independent qRT-PCR experiment comparing *Peri-15a* transcription in the intestines of Tk>*Tk*^RNAi^ flies to that in Tk> control flies. As expected, Tk>*Tk*^RNAi^ resulted in decreased transcription of *Peri- 15a* (Figure 3D).

**Figure 3:**
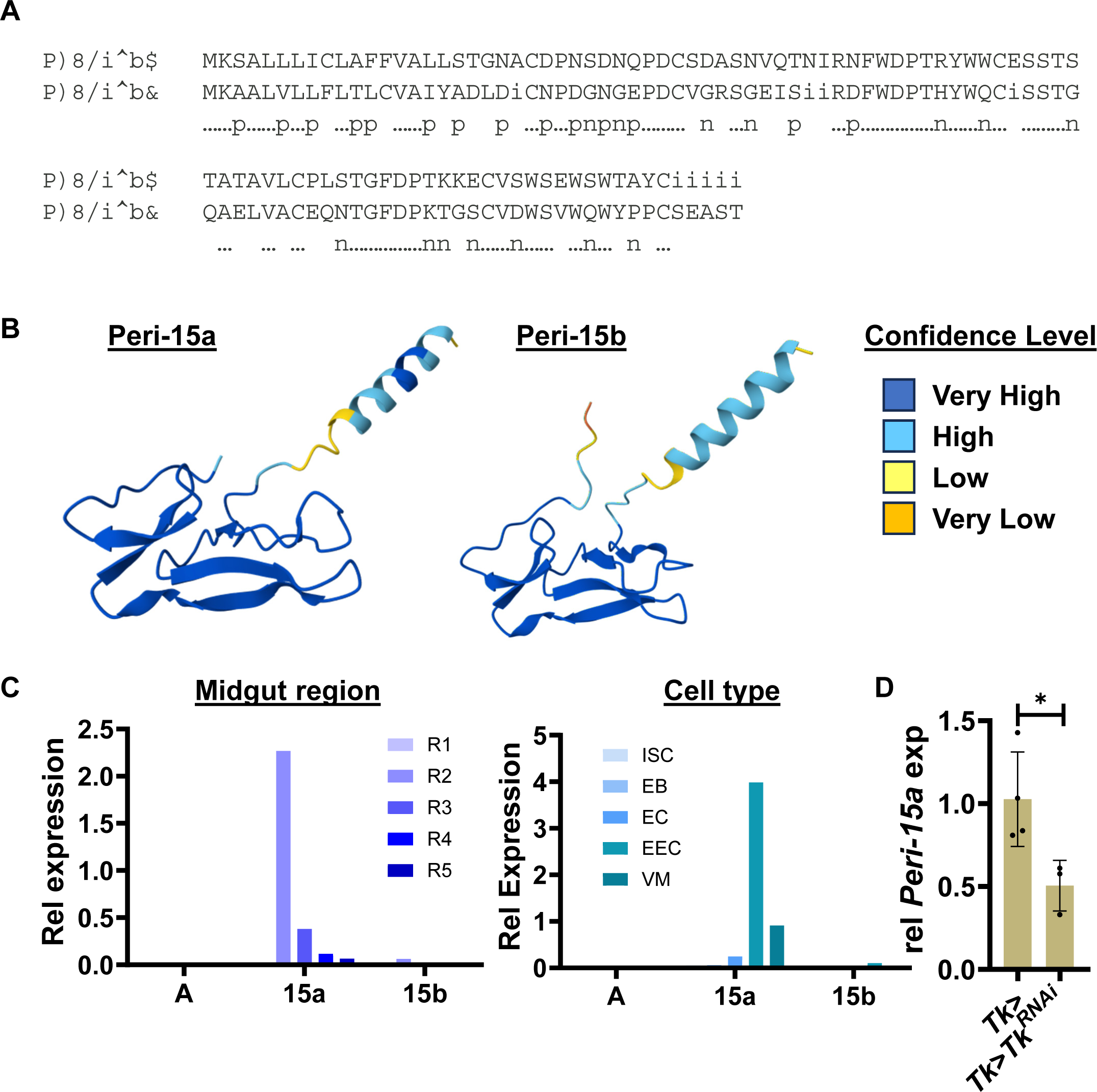
Homologs peritrophin-15a and -15b are most highly expressed in enteroendocrine cells of the *Drosophila* midgut. (A) Alignment of the amino acid sequences of Peritrophin-15a (Peri-15a) and Peritrophin-15b (Peri-15b) using Clustal Omega. The sequences are 45% identical and 58% similar. (B) Alpha fold generated structures of Peri-15a and Peri-15b. The N-terminal helices representing approximately residues 1-22 are predicted signal peptides, while residues 31-92 in Peri-15a and 38-89 in Peri-15b are predicted type 2 chitin binding domains. (C) Expression of the three midgut peritrophins, Peritrophin A, 15a, and 15b in the 5 regions of the midgut (R1-R5) and midgut cell types including stem cells (ISC), enteroblasts (EB), enterocytes (EC), enteroendocrine cells (EEC) and ventral muscle (VM). Data is from the flygutseq database (from http://flygutseq.buchonlab.com/). (D) qRT-PCR analysis of intestinal expression of Peri-15a) of the indicated flies. The mean of biological triplicates is shown. Error bars represent the standard deviation. A student’s t test was used to assess statistical significance. * p<0.05.

We next measured transcription of *Peri-15a* in the setting of *V. cholerae* infection. This significantly increased transcription of *Peri-15a* (Fig 4A). We then assessed the impact of Peri-15a on *V. cholerae* colonization. Considering that other EECs might produce Peri-15a in addition to Tk+ ones, we knocked down *Peri-15a* transcription using the pan-EEC driver Prospero (Pros>) and *Peri-15a*^RNAi^. *Peri-15a*^RNAi^ reduced *V. cholerae* colonization of the *Drosophila* intestine by approximately 100-fold (Fig 4B) and decreased intestinal *Peri-15a* transcription 2-fold but did not alter transcription of *Tk* or the gene encoding the AMP *Def* (Fig 4C and D). To further rule out an effect of Peri-15a on Tk expression, we performed Tk and lipid staining on the intestines of Pros> control and Pros>*Peri-15a*^RNAi^ flies. Pros>*Peri-15a*^RNAi^ had no impact on intestinal lipid accumulation. A minimal decrease in numbers of Tk+ cells was observed in the AMG only (Fig 4E and F). This indicates Tk is upstream of Peri-15a and that decreased colonization of the Pros>*Peri-15a*^RNAi^ intestine is not the result of increased AMP transcription. Because *V. cholerae* infection both increases transcription of *Peri-15a* and is dependent on it, we propose a model in which *V. cholerae* co-opts the host response to infection for its own benefit.

**Figure 4:**
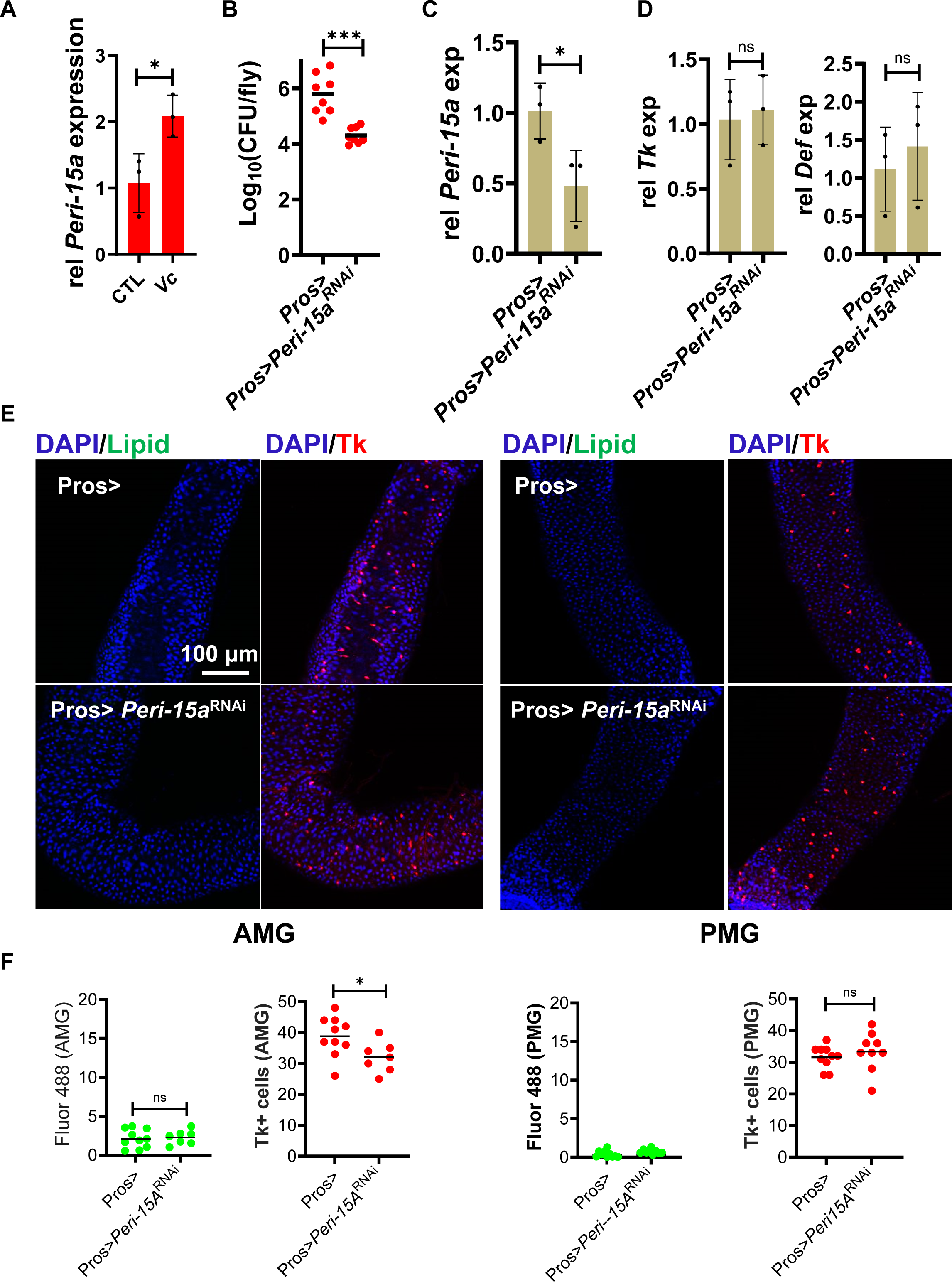
Peri-15a promotes *V. cholerae* colonization of the arthropod intestine without impacting expression of Tk or AMPs. (A) qRT-PCR analysis of *Peri-15a* transcription in the intestines of Tk> (CTL) flies fed either LB alone or inoculated with *V. cholerae* (*Vc*). The mean of biological triplicates is shown. Error bars represent the standard deviation. Significance was calculated using a student’s t test. (B) *V. cholerae* colonization of the intestines of flies with the indicated genotypes. Data were log-transformed. The mean of eight intestines is shown. Significance was calculated using a Welch’s t test. qRT-PCR analysis of transcription of (C) Peri-15 or (D) *Tk* and *Def* in the intestines of Pros> control or Pros>*Peri-15a*^RNAi^ flies. The mean of biological triplicates is shown. Error bars represent the standard deviation. Significance was calculated using a student’s t test. (E) Representative fluorescence images showing DAPI, BODIPY(Lipid), and Tk staining in the anterior midgut (AMG) and posterior midgut (PMG) of the indicated fly genotypes. Scale bars, 100 μM. (F) Quantification of (E) Representative immunofluorescence images showing DAPI and TK staining in the anterior midgut (AMG) and posterior midgut (PMG) of the indicated fly genotypes. Scale bars, 100 μM. (F) Quantification of lipid accumulation (Fluor 488) and Tk+ EECs in the AMG and PMG of the indicated fly genotypes. The mean of a minimum of seven intestines is shown. Significance was calculated using a student’s t test. ***p < 0.001, * p < 0.05, ns not significant.

Evidence from other insects suggests that Peri-15a and b do not decrease *V. cholerae* colonization by disrupting the integrity of the PM. To test the impact of altering the continuity of the PM surface on *V. cholerae* colonization, we examined intestinal chitinases. In the cotton bollworm, silencing of the *Cht4* homolog results in a less porous PM with a higher chitin content (41). *Cht4* and *Cht9*, which are the most highly expressed chitinases in the midgut, are principally expressed in enteroblasts and enterocytes of the *Drosophila* midgut (42) (Fig 5A). To determine how they impact *V. cholerae* colonization, we compared *V. cholerae* colonization of NP1>*Cht4*^RNAi^ and NP1>*Cht9*^RNAi^ flies with an NP1> enterocyte driver-only control fly. *Cht9*^RNAi^ had no effect on *V. cholerae* colonization, while *Cht4*^RNAi^ unexpectedly decreased colonization (Fig 5B). It seemed unlikely that decreased PM porosity and increased chitin content would inhibit *V. cholerae* colonization directly, since chitin has been shown to promote *V. cholerae* surface adhesion (12, 43, 44). In studies of the Tsetse fly, investigators have suggested that decreased PM porosity diminishes access of microbial products to the intestinal epithelium and, thus, reduces IMD signaling (38). Based on this precedent, we hypothesized that decreased PM porosity in NP1>*Cht4*^RNAi^ flies might decrease IMD signaling and, consequently, Tk and Peri-15A expression. To test this, we quantified transcription of *Tk*, *Def*, and *Peri-15a* in the intestines of NP1>*Cht4*^RNAi^ and NP1> flies and, at the same time, established that *Cht4*^RNAi^ decreased *Cht4* transcription (Fig 5C). In fact, all three transcripts were decreased by *Cht4*^RNAi^. This is consistent with a model in which *Cht4*^RNAi^ reduces access of microbial products to the intestinal epithelium leading to decreased IMD signaling and *Peri-15a* transcription and does not support a model in which continuity of the peritrophic matrix promotes *V. cholerae* colonization.

**Figure 5:**
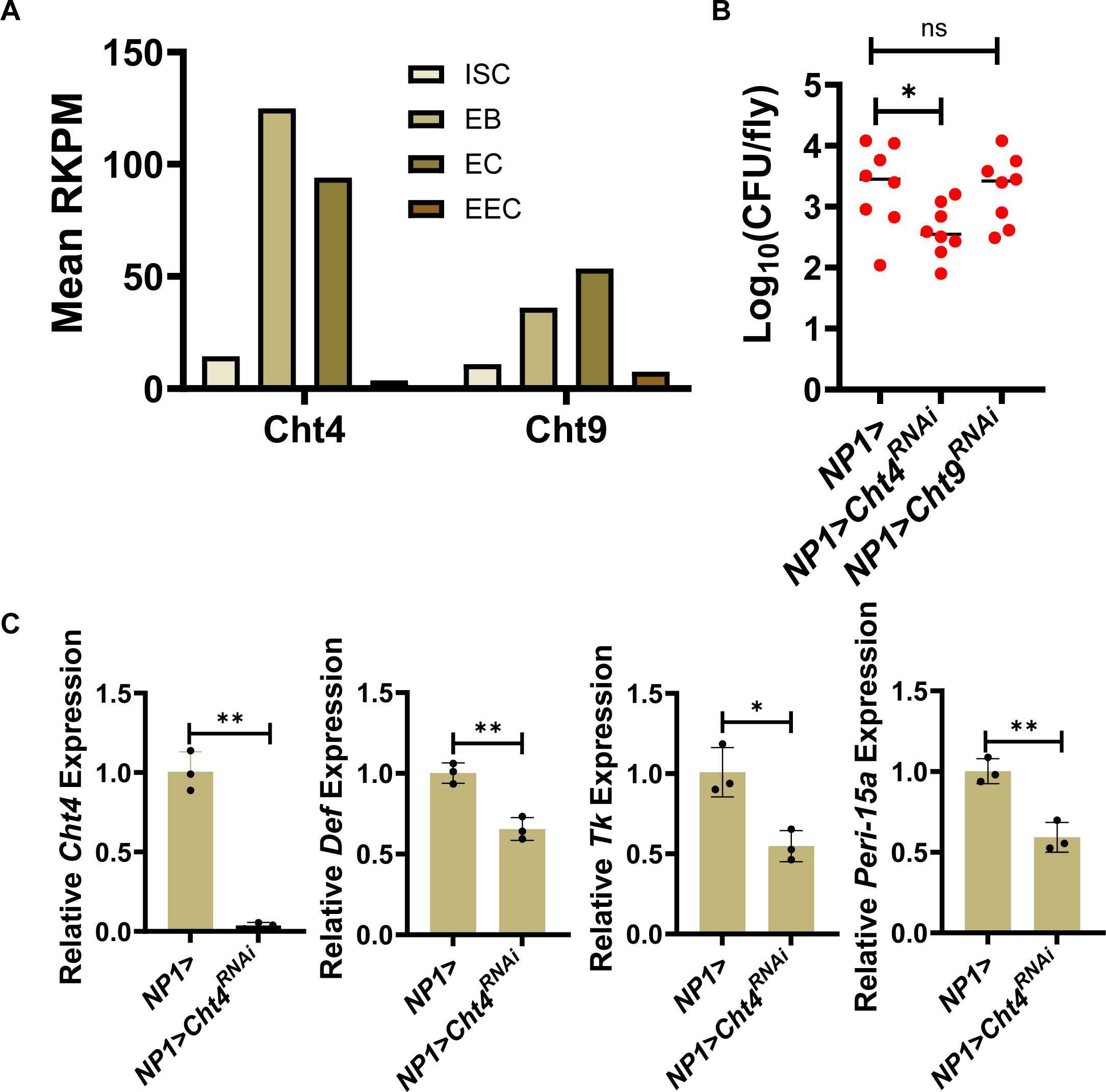
Peritrophic matrix integrity is not correlated with *V. cholerae* adhesion. (A) Expression of Chitinase 4 (Cht4) and Chitinase 9 (Cht9) in different intestinal cell types (ISC intestinal stem cells, EB enteroblasts, EC enterocytes, EEC enteroendocrine cells). database (from http://flygutseq.buchonlab.com/). (B) *V. cholerae* colonization of an enterocyte driver-only fly (NP1>) and a fly with knockdown of the indicated chitinases in enterocytes. Bars represent the mean of at least six biological replicates. An ordinary ANOVA with a Dunnett’s multiple comparisons test was performed on log_10_ transformed data to assess significance. (C) qRT-PCR measurements of *Cht4*, *Tk, Def*, and *Peri-15a* transcription in the intestines of flies of the indicated genotypes. Error bars represent the standard deviation. Significance was assessed using a Welch’s t test for *Cht4* and a student’s t test for all others. ** p<0.01, * P<0.05, ns not significant.

We have shown that decreased *Peri-15a* expression decreases *V. cholerae* colonization. We reasoned that if increased *Peri-15a* expression promoted *V. cholerae* colonization, this would serve as further evidence that a structural defect in the peritrophic matrix as a result of *Peri-15a* deficiency was not responsible for decreased *V. cholerae* colonization. *Peri-15a* expression is activated by Tk, and we have previously shown that the fly sex hormone 20-hydroxyecdysone (20E) activates Tk expression (27). Therefore, we hypothesized that intestinal *Peri-15a* expression would be activated by dietary supplementation with 20E. To test this, we compared *Tk* and *Peri-15a* transcription in the intestines of flies fed LB alone or supplemented with 20E. As shown in Figure 6A, administration of 20E to *Drosophila* increased expression of *Tk* and *Peri-15a*. Furthermore, 20E administration increased intestinal colonization by *V. cholerae* (Fig 6B). To demonstrate that this increase in colonization was dependent on Peri-15a, we assessed the impact of 20E on *V. cholerae* colonization in Pros> control and Pros>*Peri-15a*^RNAi^ flies. Supplementation with 20E increased colonization of Pros> control flies but did not significantly increase *V. cholerae* colonization in Pros>*Peri-15a* flies. This demonstrates that the increased colonization observed with 20E supplementation is dependent on Peri-15a and suggests that Peri-15a may be a host colonization factor.

**Figure 6:**
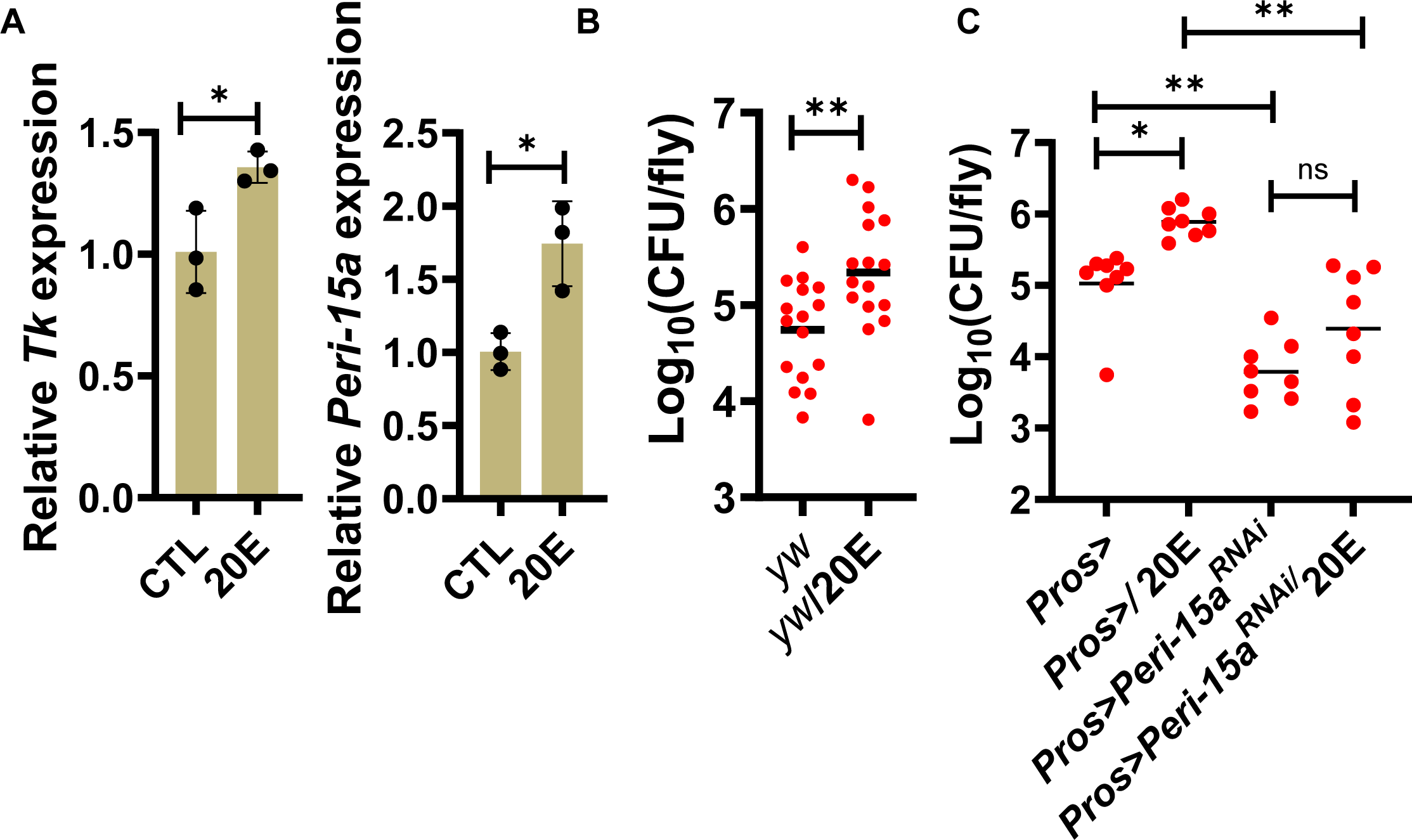
Transcriptional activation of Peritrophin-15a by 20-hydroxyecdysone increases *V. cholerae* colonization of the arthropod intestine. (A) qRT-PCR analysis of transcription of the genes tachykinin (*Tk*) and Peritrophin-15a (*Peri-15a*) in Tk> flies fed LB broth (CTL) or LB broth supplemented with 20-hydroxyecdysone (20E). The mean of biological triplicates is shown. Error bars represent the standard deviation. A student’s t-test was used to assess significance. *V. cholerae* colonization of the intestines of (B) *yw* or (C) Pros> and Pros>*Peri15a*^RNAi^ flies fed LB alone or supplemented with 20-hydroxyecdysone (20E). The mean is shown. In (B) 16 flies were used and a student’s t test was used to assess significance. In (C) at least 8 flies were used and a Welch’s ANOVA with Dunnett’s T3 multiple comparisons test was applied to assess significance. **** p<0.0001, *** p<0.001, * p<0.05, ns not significant.

We have previously shown that *V. cholerae* is able to digest the *Drosophila* peritrophic matrix (45), and *V. cholerae* can use chitin as a sole carbon and nitrogen source because its genome encodes multiple chitinases as well as transporters and enzymes that enable catabolism of chitin degradation products (12, 46–48). The latter *V. cholerae* chitin catabolic cascade is under the control of the non-canonical transcription factor ChiS (12, 47, 49). To determine the impact of chitin degradation on *V. cholerae* colonization of the arthropod gut, we assessed colonization by a *V. cholerae* Δ*chiS* mutant, which is unable to degrade chitin and utilize its metabolic products. Mutation of *chiS* improved colonization in a control fly (Fig 7A). We reasoned that this could reflect decreased damage to the peritrophic matrix leading to decreased IMD signaling and AMP expression. To test this, we compared colonization by WT *V. cholerae* and a Δ*chiS* mutant in the ΔAMP fly. The increased colonization of the Δ*chiS* mutant was preserved in the ΔAMP fly background, suggesting that this difference is independent of AMP expression. We then questioned whether improved colonization by the Δ*chiS* mutant was dependent on Tk and Peri-15a expression. As shown in Figs 7B and C, Tk>*Tk*^RNAi^ and Pros>*Peri-15a*^RNAi^ both lead to insignificant differences in colonization between WT *V. cholerae* and the Δ*chiS* mutant. Therefore, *V. cholerae* chitin catabolism interferes with the impact of Peri-15a on *V. cholerae* adhesion. We hypothesize that *V. cholerae* chitinases remove Peri-15a from the peritrophic matrix via degradation of chitin attachment sites.

**Figure 7:**
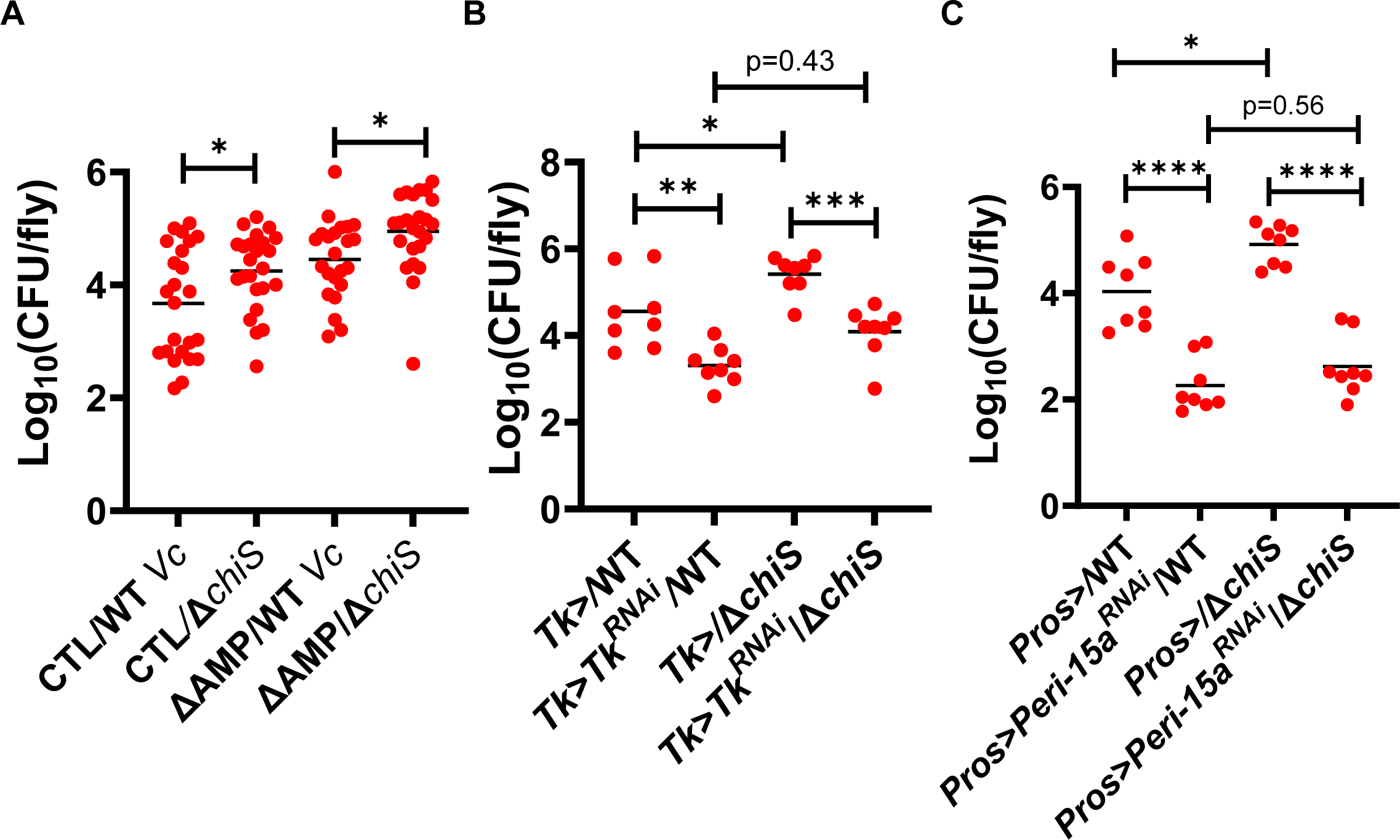
The *V. cholerae* master regulator of chitin metabolism, ChiS, decreases *V. cholerae* colonization of the arthropod intestine in a Peritrophin-15a-dependent fashion. Colonization of (A) *w^1118^*parental flies (CTL) and flies with deletion of 10 genes encoding antimicrobial peptides (ΔAMP) with either wild-type *V. cholerae* (WT) or a Δ*chiS* mutant. At least 23 flies were assessed. The mean of log_10_ transformed data is shown. A one-way ordinary ANOVA was used to assess significance. Colonization of (B) Tk> driver only and Tk>*Tk*^RNAi^ flies or (C) Pros> driver-only and Pros>*Peri15a*^RNAi^ flies with the indicated *V. cholerae* strains. Eight flies were quantified for each condition. The mean of log_10_ transformed data is shown. An ordinary one-way ANOVA with Dunnett’s multiple comparisons test was used to assess significance. The mean of biological triplicates is shown. Error bars represent the standard deviation. A student’s t test was used to assess significance. **** p,0.0001, ** p<0.01, * p<0.05, ns not significant.

*V. cholerae* has been associated with zooplankton in aquatic environments, and there is evidence that *V. cholerae* colonizes these organisms using specific adhesins (5, 50–55). Furthermore, transcription of copepod genes encoding innate immune effectors and chitin binding proteins, including a homolog of Peri-15a, were increased in response to *Vibrio* colonization (32). To understand the relevance of our findings to the natural environment of *V. cholerae*, we searched for structural homologs of Peri-15a in the AlphaFold protein structure database. Highly similar Peri-15a homologs were present in diverse insects including mosquitos, beetles, butterflies, and sand flies (Fig 8). Structural homologs of Peri-15a were also identified in the genomes of marine non-biting midges, rotifers, marine copepods, zooplankton, and fungi, all of which are found in the marine aquatic environments inhabited by *V. cholerae*. Taken together, these results suggest that Peri-15a homologs are present in natural environmental hosts of *V. cholerae* and that intestinal colonization with *V. cholerae* increases expression of these proteins by activating the intestinal innate immune response.

**Figure 8:**
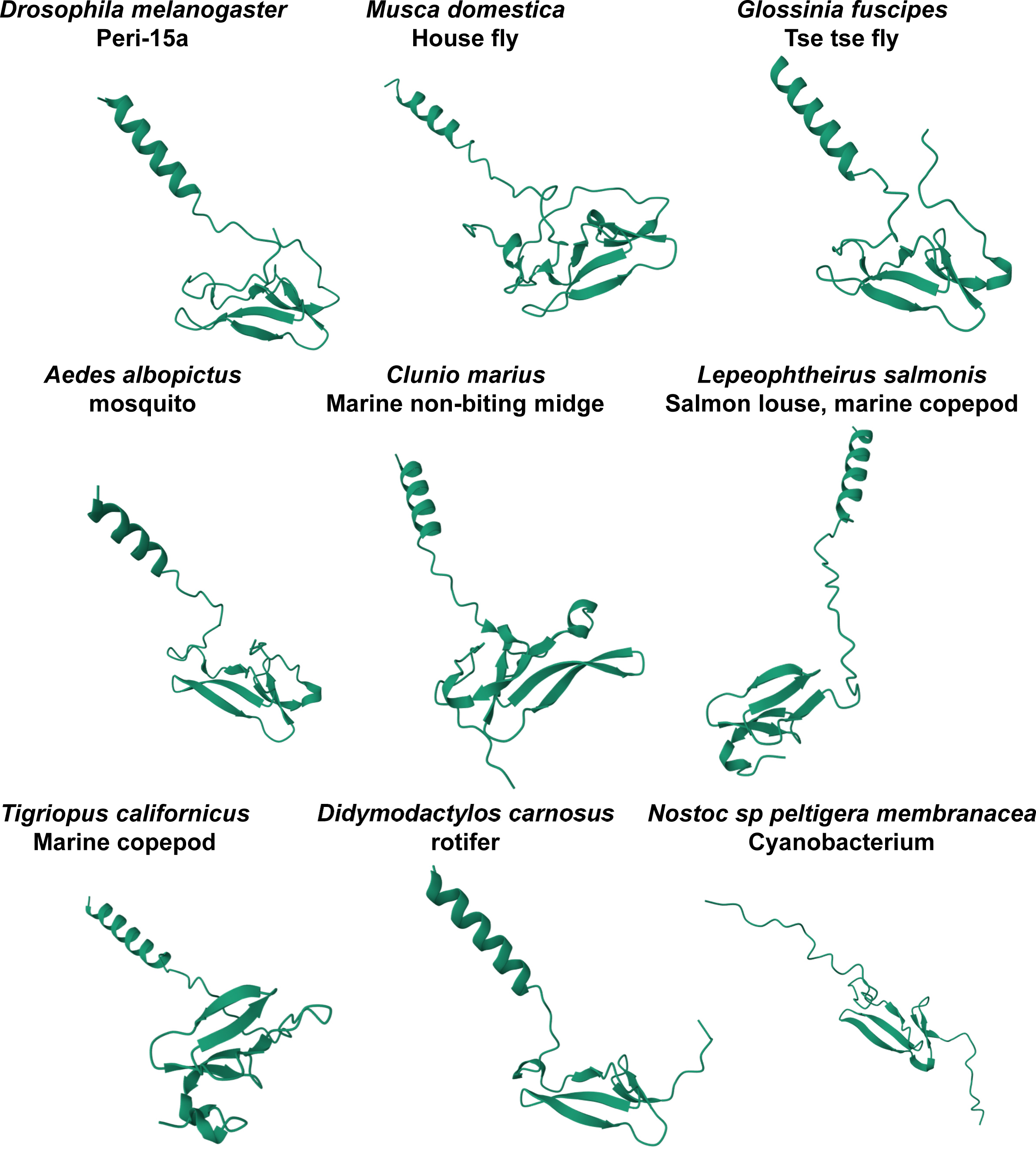
Structural homologs of Peri-15a are found in cyanobacteria, fungi, rotifers, copepods and marine insects. Alpha fold AFDB50, AFDB Fold, and AFDB50MMseq2 were used to find structural homologs of *Drosophila* Peri-15a. The structures of homologs found in insect vectors, copepods, a rotifer, and a cyanobacterium are shown.

## Discussion

Arthropods figure prominently in the environmental life cycle of *V. cholerae*. Here we show that a small, widely conserved arthropod protein consisting of a secretion peptide and a single type 2 chitin binding domain promotes *V. cholerae* colonization of the arthropod intestine. This protein, termed Peri-15a in the model arthropod *Drosophila melanogaster*, is regulated by the enteroendocrine peptide Tk, expressed almost exclusively in enteroendocrine cells of the adult midgut and secreted and incorporated into the PM. Homologs of Peri-15a are widely distributed in terrestrial arthropods and zooplankton that have been associated with *V. cholerae.* While the goal of increasing Peri-15a expression may be to protect the PM from microbial degradation, we propose that *V. cholerae* has co-opted Peri-15a as an intestinal colonization factor of environmental arthropod hosts.

Environmental studies suggest that the presence of *V. cholerae* in the water column is correlated with zooplankton such as cladocerans, copepods, and rotifers (5, 56, 57). Attachment to the exoskeletons of zooplankton has also been observed in the laboratory (10, 51, 54). Arthropods, which include cladocerans and copepods, have been found to dominate the zooplankton population in diverse environments (58–62). In the laboratory, some have observed that *V. cholerae* attaches much more abundantly to dead zooplankton arthropods or their molts as compared with live ones (52, 55). This is not surprising as the arthropod exoskeleton is designed to be an impermeable, microbe-resistant barrier (63). The outermost layer is comprised of hydrophobic lipoproteins, while inner layers consist of complex bundles of protein-chitin fibers. Thus, shed exoskeletons provide more ready access to chitin. In contrast, the intestine of arthropods provides a more protected environment for microbes such as *V. cholerae*, which have been found in the intestines of zooplankton (22, 64). While the exoskeleton is impermeable, the PM of the midgut is designed for permeability as it must allow transit of nutrients and small molecules (65). The chitin fibers are thin and more readily accessible. Here we provide evidence for a specific interaction between *V. cholerae* and a protein incorporated into the arthropod PM.

PM porosity has been shown to modulate the burden of intestinal bacteria in other insects. PM porosity is critical because it must balance permeability to nutrients, which is essential, with permeability to microbe or microbe metabolism-associated molecular products, which may lead to chronic immune stimulation and intestinal damage. Consistent with this model, *Cht4* knockdown in the insect *Helicoverpa armigera* decreased PM porosity and body weight (41). Furthermore, in the tsetse fly, investigators found that chitin synthase knockdown activated the intestinal innate immune response presumably by increasing PM permeabilty. This reduced the intestinal burden of orally administered Gram-negative pathogens such as *Serratia marcescens* and *Enterobacter* sp (38).

Large PM proteins have multiple chitin binding domains that are thought to crosslink single chitin fibers to each other to form fibrils that provide structural rigidity to the matrix (25). In contrast, Peri-15a, which is most highly expressed in enteroendocrine cells of the AMG, consists of only one chitin binding domain. Biochemical studies have shown that Peri-15a binds chitin directly at very low stoichiometry *in vitro* and is tightly associated with the PM (66). Therefore, rather than crosslinking chitin fibers, it has been suggested that Peri-15a is a capping protein that protects single chitin strands from exochitinases. Homologs of Peri-15a and b, known as Proventriculin 1 and 2, have been identified and studied in the PM of the tsetse fly (67). Consistent with the model of Peri-15a as a capping protein, these investigators found that knockdown of these proteins left the permeability of the peritrophic membrane largely intact (38). Therefore, it is unlikely that Peri-15a promotes *V. cholerae* colonization through a major structural rearrangement of the PM.

Here, we report that *Cht4*^RNAi^, which is predicted to reduce matrix porosity, decreased the intestinal innate immune response, expression of *Peri-15a*, and intestinal colonization by *V. cholerae*. In contrast to arthropod *Cht4*^RNAi^, deletion of *V. cholerae chiS*, the master regulator of *Vibrio* chitin catabolism increased colonization of the arthropod intestine (12, 49, 68). ChiS activates transcription of at least four of the five proven *Vibrio* endochitinases: ChiA1 (VC1952), ChiA2 (VCA0027), VC0769, and VCA0700 (11, 12), and we have previously presented evidence that *V. cholerae* thins the chitinous layer of the *Drosophila* peritrophic matrix, suggesting that these chitinases are active in the *Drosophila* intestine (45). Knockdown of the arthropod chitinases decreases the porosity of the peritrophic matrix and expression of Peri-15a. In contrast, *V. cholerae* chitinases act on naturally formed peritrophic matrices to digest them, likely removing Peri-15A at the same time. It has previously been proposed that *V. cholerae* attaches to and degrades environmental chitin to release nutrients required for its growth (9). Our results suggest that degradation of chitin by *V. cholerae* promotes detachment from the chitin-rich PM. Taken together, these findings support a model in which Peri-15a does not enhance *V. cholerae* intestinal colonization by altering peritrophic matrix porosity. Rather, we propose that Peri-15a either binds a *V. cholerae* adhesin or reveals a binding site in the PM for such an adhesin.

GbpA (VCA0811) and FrhA (VC1620) are two *V. cholerae* adhesins that mediate binding to abiotic surfaces, plankton and the mammalian intestinal epithelium(50, 51). GbpA is a secreted, four-domain 485aa protein whose expression is increased by chitin but which, unlike the *V. cholerae* chitinases, is not under ChiS control (12, 69). Domains 1 and 4 mediate chitin binding, domain 1 also mediates binding to mammalian mucins, and domains 2 and 3 interact with the *V. cholerae* surface, enabling GbpA to form bridges between the bacterium and other surfaces (69). While biochemical studies have shown that GbpA can function as a chitinase, there is no *in vivo* evidence for this as of yet (11, 70). FrhA is a secreted 2,251 amino acid protein first identified because it was co-regulated with the flagellar apparatus (71). It contains three large N-terminal Calx-beta Ca^2+^ binding domains and 4 C-terminal cadherin-like Ca^2+^ binding domains. Amino acids residues 1155-1350 were recently identified as a peptide-binding domain that is sufficient to enhance binding to abiotic surfaces, epithelial cells, the mouse intestine, and diatoms (51). Either of these could serve as the adhesin mediating attachment to the PM via Peri-15a.

Environmental studies suggest that the presence of zooplankton and *V. cholerae* are correlated in the aquatic environment (5, 56, 57). Furthermore, the observation that *V. cholerae* possesses all the transporters and enzymes required to utilize chitin as a sole energy source, that the *V. cholerae* genome encodes a transcription factor dedicated to regulating its interaction with chitins, and that *V. cholerae* becomes naturally competent in the presence of chitin suggest that chitinous surfaces play an important role in the *V. cholerae* life cycle (12, 13, 47). Here we provide further evidence for a specific interaction between *V. cholerae* and the arthropod intestinal PM by identifying a protein component of the PM that does not significantly alter PM structure but, nevertheless, enhances *V. cholerae* colonization.

## Experimental Methods

### *Drosophila* husbandry and strains

All fly stocks used in this study were raised on standard fly food containing 16.5 g/L yeast, 9.5 g/L soy flour, 71 g/L cornmeal, 5.5 g/L agar, 5.5 g/L malt, 7.5% corn syrup, and 0.4% propionic acid in a humidified 12 hr day-night cycle incubator at 25°C. Females flies (4-7 day old) were used in all experiments. The following RNAi and driver fly stocks were obtained from the Bloomington Drosophila Stock Center (BDSC): *Tk*^RNAi^ (BL2500), *Rel^RNAi^* (BL28943)*, TkR99D^RNAi^*(BL27513)*, Cht4*^RNAi^ (BL65001), *Peritrophin-15a*^RNAi^ (BL55987), and Pros-Gal4/Sb (BL84276). Control lines TRiP (BL36303) and *y*^1^*w*^1^ (BL1495) were also obtained from the BDSC. The Tk-Gal4 and NP1-Gal4 driver lines were a kind gift from the Veenstra and Perrimon laboratories (31, 72). The AMP-deficient flies and the corresponding parental strain were generously provided by the Lemaitre laboratory (34).

Bacterial strains and media. The wild-type *V. cholerae* strain and the parental background for all mutants was quorum sensing-competent *V. cholerae* 01 El Tor strain C6706 (C6706 str 2), a kind gift of the late Ronald Taylor. The Δ*chiS* mutant was constructed by double homologous recombination as previously described using the primers listed in Table S2 (18). *V. cholerae* was cultured at 27°C in LB broth (Miller formulation, Difco) or on LB agar (Difco) supplemented with 100 µg/mL streptomycin. Where noted, 20-hydroxyecdysone was added to LB broth for *Drosophila* colonization studies (10 µM, Sigma).

*V. cholerae* mutant construction. An in-frame *V. cholerae* Δ*chiS* mutant was constructed by double homologous recombination as previously described (73). ChiS is comprised of 1177 amino acids. In this mutant, residues 172 to 1053 were deleted including all conserved domains.

*V. cholerae* colonization assays. For colonization assays, *V. cholerae* was cultured overnight in LB broth and then diluted 10-fold in fresh LB broth. 3 mL of this suspension was infused into autoclaved cellulose plugs inserted at the bottom of fly vials. 4–7-day old flies were placed in these prepared vials for 48 hours at 25 °C and then transferred to fresh vials containing cellulose plugs infused with 3 mL sterile solution of phosphate-buffered saline for a 24-hour to washout non-adherent bacteria. Individual flies treated in this manner were then homogenized in 200 µL of 1x PBS using a TissueLyser III system (Qiagen). Serial 10-fold dilutions were prepared, and 20 μl of these dilutions were plated on LB agar plates supplemented with 100 µg/mL streptomycin. The plates were incubated at 27°C for 24 hours and colony-forming units (CFU) in each dilution were enumerated. This was used to calculate *V. cholerae* CFU/fly.

*Drosophila* RNA extraction, RNAseq, and RT-qPCR. Total RNA was isolated from 10-15 fly intestines using TRIzol reagent (Thermo Fisher Scientific 15596026) and the Direct-zol RNA Miniprep plus kit (Zymo Research R2070). Three to six biological replicates per condition/genotype were performed. For RNA sequencing analysis, the RNA was submitted to the Molecular Biology Core Facilities (MBCF) at the Dana Farber Cancer Institute (DFCI) for next-generation sequencing (NGS) library preparation, sequencing, and analysis (https://mbcf.dana-farber.org/totalrnaseq.html). Roche Kapa mRNA HyperPrep strand specific sample preparation kits were used to prepare libraries on a Beckman Coulter Biomek i7 from 200ng of purified total RNA. The resulting dsDNA libraries were quantified by Qubit fluorometer and Agilent TapeStation 4200. Uniquely dual indexed libraries were pooled in an equimolar ratio and shallowly sequenced on an Illumina MiSeq to further evaluate library quality and pool balance. The final pool was sequenced on an Illumina NovaSeq X Plus targeting 40 million 150bp read pairs per library. Sequenced reads were aligned to the UCSC hg38 reference genome assembly and gene counts were quantified using STAR (v2.7.3a) and Salmon (74, 75). Differential gene expression testing was performed by DESeq2 (v1.22.1) (76). RNAseq analysis was performed using the VIPER snakemake pipeline (77).

For qRT-PCR, the iScript™ cDNA synthesis kit (Bio-Rad 1708891) was used to synthesize cDNA from 500ng of total RNA. The iTaq™ Universal SYBR^®^ Green Supermix (Bio-rad 1725121) was used to perform qPCR of target gene transcripts on a QuantStudio™ Real-Time PCR system 3 (Thermo Fisher Scientific). Quantification cycle values (Cq) were obtained and used to calculate target gene transcription normalized to Actin. Primers used in this study are listed in Table S2.

Lipid staining and immunofluorescence of *Drosophila* intestine. Fly intestines were dissected in 1x PBS and fixed in a 4% paraformaldehyde (PFA), 0.1% PBS-Tween 20 (PBT) solution for 20 minutes. For lipid staining only, the intestines were then washed three times for 10 minutes with PBT, followed by incubation in staining solution #1 (PBT + 1:1000 DAPI (Invitrogen D1306) + 1:500 BODIPY 493/503 (Invitrogen D3922)) for 2 hours. For lipid staining and Tk-immunofluorescence experiments, intestines were washed three times for 10 minutes in PBT and then left in blocking solution (PBT + 0.1% Triton X-100 (Sigma-Aldrich 9002-93-1) + 2% BSA (Sigma-Aldrich 9048-46-8)) for 1 hour. After this blocking step, intestines were left in Rabbit anti-Tk primary antibody solution (blocking solution + 1:500 anti-Tk) overnight at 4°C. The next day the guts were washed three times with PBT for 10 minutes, followed by incubation in staining solution #2 (blocking solution + 1:1000 DAPI (Invitrogen D1306) + 1:500 BODIPY 493/503 (Invitrogen D3922) + 1:500 Alexa 594-conjugated Goat anti-rabbit secondary antibody (Thermo Fisher Scientific A11012)) for 2 hours. After incubation in the appropriate staining solution, intestines were washed three times for 10-minutes in PBT and then mounted in Vectashield® antifade mounting medium (Vector Laboratories H-1000-10). Intestines were imaged using a Zeiss LSM 980 confocal microscope using a 40X (Fig 2) or 25X (Fig 4) oil objective. Unless otherwise noted, all steps were done at room temperature. Lipid accumulation was quantified using ImageJ (FIJI) to measure total BODIPY fluorescence. The corrected total cell fluorescence (CTCF) was calculated by dividing total BODIPY fluorescence by the total area and subtracting background fluorescence from this measurement. Tk-expressing cells were quantified manually. A minimum of six fly intestines per genotype were evaluated for quantification of Tk-expressing cells and BODIPY fluorescence.

Statistical analysis. All data were graphed and analyzed using Prism 10.0 software (GraphPad). Measurements shown in each graph represent the mean values of at least three biological replicates for qRT-PCR, at least six flies for colonization experiments, and at least six intestines for microscopy. The mean is shown for all experiments, and error bars represent the standard deviation. Where noted, data were log-transformed to obtain a normal distribution. Appropriate statistical tests were chosen based on numbers of conditions assessed and tests of normality and variance. The statistical test used in each graph is specified in the figure legend.

## Data availability statement

RNAseq data have been deposited in the NCBI repository (accession no. GSE294931).

## Acknowledgements

This work was supported by NIH R01AI158247 and NIH R01AI162701 to P.I.W. and NIH F31DK130254 to D. B. Anti-TK antibodies were generously provided by Jan Veenstra and AMP-deficient flies and the corresponding parental strain were generously provided by Bruno Lemaitre. The TK-Gal4 and NP1-Gal4 (Myo1A-Gal4) driver flies were kind gifts from Norbert Perrimon. Stocks obtained from the Bloomington Drosophila Stock Center (NIH P40OD018537) were used in this study. Microscopy images were acquired at the Microscopy Resources on the North Quad (MicRoN) core at Harvard Medical School. We thank Paola Montero Lopis at the MicRoN core for providing expertise with image acquisition and quantification.

## Supplementary Materials

**Table S1: Genes differentially regulated in the intestines of uninfected Tk>*Tk*^RNAi^ flies as compared with Tk> driver only flies (Tk>*Tk*^RNAi^/Tk>). Driver-only flies were crossed to a Trip control fly line (y sc v).**

**Table S2: Sequences of primers used for qRT-PCR.**

## References

1. A. Hsiao, J. Zhu, Pathogenicity and virulence regulation of Vibrio cholerae at the interface of host-gut microbiome interactions. Virulence 11, 1582–1599 (2020).

2. S. I. Miyoshi et al., Isolation of Vibrio cholerae and Vibrio vulnificus from Estuarine Waters, and Genotyping of V. vulnificus Isolates Using Loop-Mediated Isothermal Amplification. Microorganisms 12 (2024).

3. S. K. Banerjee, R. Rutley, J. Bussey, Diversity and Dynamics of the Canadian Coastal Vibrio Community: an Emerging Trend Detected in the Temperate Regions. J Bacteriol 200 (2018).

4. M. L. Lizarraga-Partida et al., Association of Vibrio cholerae with plankton in coastal areas of Mexico. Environ Microbiol 11, 201–208 (2009).

5. G. C. de Magny et al., Role of Zooplankton Diversity in Vibrio cholerae Population Dynamics and in the Incidence of Cholera in the Bangladesh Sundarbans. Appl Environ Microbiol 77, 6125–6132 (2011).

6. M. Broza, H. Gancz, Y. Kashi, The association between non-biting midges and Vibrio cholerae. Environ Microbiol 10, 3193–3200 (2008).

7. R. Sela, B. K. Hammer, M. Halpern, Quorum-sensing signaling by chironomid egg masses’ microbiota, affects haemagglutinin/protease (HAP) production by Vibrio cholerae. Mol Ecol 30, 1736–1746 (2021).

8. A. Huq et al., Ecological relationships between *Vibrio cholerae* and planktonic crustacean copepods. Appl Environ Microbiol 45, 275–283 (1983).

9. C. Pruzzo, L. Vezzulli, R. R. Colwell, Global impact of Vibrio cholerae interactions with chitin. Environ Microbiol 10, 1400–1410 (2008).

10. D. A. Chiavelli, J. W. Marsh, R. K. Taylor, The mannose-sensitive hemagglutinin of *Vibrio cholerae* promotes adherence to zooplankton. Appl Environ Microbiol 67, 3220–3225. (2001).

11. C. A. Hayes, T. N. Dalia, A. B. Dalia, Systematic genetic dissection of chitin degradation and uptake in Vibrio cholerae. Environ Microbiol 19, 4154–4163 (2017).

12. K. L. Meibom et al., The Vibrio cholerae chitin utilization program. Proc Natl Acad Sci U S A 101, 2524–2529 (2004).

13. K. L. Meibom, M. Blokesch, N. A. Dolganov, C. Y. Wu, G. K. Schoolnik, Chitin induces natural competence in Vibrio cholerae. Science 310, 1824–1827 (2005).

14. M. Blokesch, G. K. Schoolnik, Serogroup conversion of Vibrio cholerae in aquatic reservoirs. PLoS Pathog 3, e81 (2007).

15. L. Kamareddine et al., Activation of Vibrio cholerae quorum sensing promotes survival of an arthropod host. Nat Microbiol 3, 243–252 (2018).

16. Z. Wang, S. Hang, A. E. Purdy, P. I. Watnick, Mutations in the IMD pathway and mustard counter Vibrio cholerae suppression of intestinal stem cell division in Drosophila. MBio 4, e00337–00313 (2013).

17. A. E. Purdy, P. I. Watnick, Spatially selective colonization of the arthropod intestine through activation of Vibrio cholerae biofilm formation. Proc Natl Acad Sci U S A 108, 19737–19742 (2011).

18. A. S. Vanhove et al., Vibrio cholerae ensures function of host proteins required for virulence through consumption of luminal methionine sulfoxide. PLoS Pathog 13, e1006428 (2017).

19. B. E. Jugder et al., Vibrio cholerae high cell density quorum sensing activates the host intestinal innate immune response. Cell Rep 40, 111368 (2022).

20. N. S. Blow et al., Vibrio cholerae infection of Drosophila melanogaster mimics the human disease cholera. PLoS Pathog 1, e8 (2005).

21. D. Fast, B. Kostiuk, E. Foley, S. Pukatzki, Commensal pathogen competition impacts host viability. Proc Natl Acad Sci U S A 115, 7099–7104 (2018).

22. T. Xu, A. Novotny, S. Zamora-Terol, P. A. Hamback, M. Winder, Dynamics of Gut Bacteria Across Different Zooplankton Genera in the Baltic Sea. Microb Ecol 87, 48 (2024).

23. D. Fast et al., Vibrio cholerae-Symbiont Interactions Inhibit Intestinal Repair in Drosophila. Cell Rep 30, 1088–1100 e1085 (2020).

24. A. C. Wong, A. S. Vanhove, P. I. Watnick, The interplay between intestinal bacteria and host metabolism in health and disease: lessons from Drosophila melanogaster. Dis Model Mech 9, 271–281 (2016).

25. S. Muthukrishnan, H. Merzendorfer, Y. Arakane, Q. Yang, Chitin Organizing and Modifying Enzymes and Proteins Involved In Remodeling of the Insect Cuticle. Adv Exp Med Biol 1142, 83–114 (2019).

26. P. I. Watnick, B. E. Jugder, Microbial Control of Intestinal Homeostasis via Enteroendocrine Cell Innate Immune Signaling. Trends Microbiol 28, 141–149 (2020).

27. B. E. Jugder, L. Kamareddine, P. I. Watnick, Microbiota-derived acetate activates intestinal innate immunity via the Tip60 histone acetyltransferase complex. Immunity 54, 1683–1697 e1683 (2021).

28. L. Kamareddine, W. P. Robins, C. D. Berkey, J. J. Mekalanos, P. I. Watnick, The Drosophila Immune Deficiency Pathway Modulates Enteroendocrine Function and Host Metabolism. Cell Metab 28, 449–462 e445 (2018).

29. D. Dutta et al., Regional Cell-Specific Transcriptome Mapping Reveals Regulatory Complexity in the Adult Drosophila Midgut. Cell Rep 12, 346–358 (2015).

30. A. Kleino, N. Silverman, The Drosophila IMD pathway in the activation of the humoral immune response. Dev Comp Immunol 42, 25–35 (2014).

31. W. Song, J. A. Veenstra, N. Perrimon, Control of lipid metabolism by tachykinin in Drosophila. Cell Rep 9, 40–47 (2014).

32. A. A. Almada, A. M. Tarrant, Vibrio elicits targeted transcriptional responses from copepod hosts. FEMS Microbiol Ecol 92, fiw072 (2016).

33. M. Varadi et al., AlphaFold Protein Structure Database in 2024: providing structure coverage for over 214 million protein sequences. Nucleic Acids Res 52, D368–D375 (2024).

34. M. A. Hanson et al., Synergy and remarkable specificity of antimicrobial peptides in vivo using a systematic knockout approach. Elife 8 (2019).

35. A. Ozturk-Colak et al., FlyBase: updates to the Drosophila genes and genomes database. Genetics 227 (2024).

36. D. Hegedus, M. Erlandson, C. Gillott, U. Toprak, New insights into peritrophic matrix synthesis, architecture, and function. Annu Rev Entomol 54, 285–302 (2009).

37. T. Kuraishi, O. Binggeli, O. Opota, N. Buchon, B. Lemaitre, Genetic evidence for a protective role of the peritrophic matrix against intestinal bacterial infection in Drosophila melanogaster. Proc Natl Acad Sci U S A 108, 15966–15971 (2011).

38. B. L. Weiss, A. F. Savage, B. C. Griffith, Y. Wu, S. Aksoy, The peritrophic matrix mediates differential infection outcomes in the tsetse fly gut following challenge with commensal, pathogenic, and parasitic microbes. J Immunol 193, 773–782 (2014).

39. F. Madeira et al., The EMBL-EBI Job Dispatcher sequence analysis tools framework in 2024. Nucleic Acids Res 52, W521–W525 (2024).

40. J. Jumper et al., Highly accurate protein structure prediction with AlphaFold. Nature 596, 583–589 (2021).

41. D. Q. Hu et al., The effect of group IV chitinase, HaCHT4, on the chitin content of the peritrophic matrix (PM) during larval growth and development of Helicoverpa armigera. Pest Manag Sci 78, 1815–1823 (2022).

42. D. P. Leader, S. A. Krause, A. Pandit, S. A. Davies, J. A. T. Dow, FlyAtlas 2: a new version of the Drosophila melanogaster expression atlas with RNA-Seq, miRNA-Seq and sex-specific data. Nucleic Acids Res 46, D809–D815 (2018).

43. S. Sun, Q. X. Tay, S. Kjelleberg, S. A. Rice, D. McDougald, Quorum sensing-regulated chitin metabolism provides grazing resistance to Vibrio cholerae biofilms. ISME J 9, 1812–1820 (2015).

44. B. R. Wucher et al., Vibrio cholerae filamentation promotes chitin surface attachment at the expense of competition in biofilms. Proc Natl Acad Sci U S A 116, 14216–14221 (2019).

45. S. Hang et al., The acetate switch of an intestinal pathogen disrupts host insulin signaling and lipid metabolism. Cell Host Microbe 16, 592–604 (2014).

46. S. Nahar et al., Role of Shrimp Chitin in the Ecology of Toxigenic Vibrio cholerae and Cholera Transmission. Front Microbiol 2, 260 (2011).

47. X. Li, S. Roseman, The chitinolytic cascade in Vibrios is regulated by chitin oligosaccharides and a two-component chitin catabolic sensor/kinase. Proc Natl Acad Sci U S A 101, 627–631 (2004).

48. E. Y. Markov, E. S. Kulikalova, L. Y. Urbanovich, V. S. Vishnyakov, S. V. Balakhonov, Chitin and Products of Its Hydrolysis in Vibrio cholerae Ecology. Biochemistry (Mosc) 80, 1109–1116 (2015).

49. C. A. Klancher, S. Yamamoto, T. N. Dalia, A. B. Dalia, ChiS is a noncanonical DNA-binding hybrid sensor kinase that directly regulates the chitin utilization program in Vibrio cholerae. Proc Natl Acad Sci U S A 117, 20180–20189 (2020).

50. T. J. Kirn, B. A. Jude, R. K. Taylor, A colonization factor links Vibrio cholerae environmental survival and human infection. Nature 438, 863–866 (2005).

51. C. J. Lloyd et al., A peptide-binding domain shared with an Antarctic bacterium facilitates Vibrio cholerae human cell binding and intestinal colonization. Proc Natl Acad Sci U S A 120, e2308238120 (2023).

52. M. L. Tamplin, A. L. Gauzens, A. Huq, D. A. Sack, R. R. Colwell, Attachment of *Vibrio cholerae* serogroup O1 to zooplankton and phytoplankton of Bangladesh waters. Appl. Environ. Microbiol. 56, 1977–1980 (1990).

53. B. N. Shukla, D. V. Singh, S. C. Sanyal, Attachment of non-culturable toxigenic *Vibrio cholerae* O1 and non-O1 and *Aeromonas* spp. to the aquatic arthropod *Gerris spinolae* and plants in the River Ganga, Varanasi. FEMS Immunol. and Medical Microbiol. 12, 113–120 (1995).

54. T. K. Rawlings, G. M. Ruiz, R. R. Colwell, Association of Vibrio cholerae O1 El Tor and O139 Bengal with the Copepods Acartia tonsa and Eurytemora affinis. Appl Environ Microbiol 73, 7926–7933 (2007).

55. R. S. Mueller et al., Vibrio cholerae strains possess multiple strategies for abiotic and biotic surface colonization. J Bacteriol 189, 5348–5360 (2007).

56. S. Schauer et al., Dynamics of Vibrio cholerae abundance in Austrian saline lakes, assessed with quantitative solid-phase cytometry. Environ Microbiol 17, 4366–4378 (2015).

57. J. W. Turner, L. Malayil, D. Guadagnoli, D. Cole, E. K. Lipp, Detection of Vibrio parahaemolyticus, Vibrio vulnificus and Vibrio cholerae with respect to seasonal fluctuations in temperature and plankton abundance. Environ Microbiol 16, 1019–1028 (2014).

58. Y. G. Wang, L. C. Tseng, M. Lin, J. S. Hwang, Vertical and geographic distribution of copepod communities at late summer in the Amerasian Basin, Arctic Ocean. PLoS One 14, e0219319 (2019).

59. M. Gluchowska et al., Variations in the structural and functional diversity of zooplankton over vertical and horizontal environmental gradients en route to the Arctic Ocean through the Fram Strait. PLoS One 12, e0171715 (2017).

60. Z. Zhang et al., Spatio-Temporal Variations of Zooplankton and Correlations with Environmental Parameters around Tiaowei Island, Fujian, China. Int J Environ Res Public Health 19 (2022).

61. V. Venkataramana, L. Gawade, M. D. Bharathi, V. Sarma, Role of salinity on zooplankton assemblages in the tropical Indian estuaries during post monsoon. Mar Pollut Bull 190, 114816 (2023).

62. G. Boldrocchi, Y. Moussa Omar, D. Rowat, R. Bettinetti, First results on zooplankton community composition and contamination by some persistent organic pollutants in the Gulf of Tadjoura (Djibouti). Sci Total Environ 627, 812–821 (2018).

63. P. Mrak, U. Bogataj, J. Strus, N. Znidarsic, Cuticle morphogenesis in crustacean embryonic and postembryonic stages. Arthropod Struct Dev 46, 77–95 (2017).

64. M. R. Sochard, D. F. Wilson, B. Austin, R. R. Colwell, Bacteria associated with the surface and gut of marine copepods. Appl Environ Microbiol 37, 750–759 (1979).

65. B. Moussian, The apical plasma membrane of chitin-synthesizing epithelia. Insect Sci 20, 139–146 (2013).

66. G. Wijffels et al., A novel family of chitin-binding proteins from insect type 2 peritrophic matrix. cDNA sequences, chitin binding activity, and cellular localization. J Biol Chem 276, 15527–15536 (2001).

67. C. Rose et al., An investigation into the protein composition of the teneral Glossina morsitans morsitans peritrophic matrix. PLoS Negl Trop Dis 8, e2691 (2014).

68. C. A. Klancher et al., The ChiS-Family DNA-Binding Domain Contains a Cryptic Helix-Turn-Helix Variant. mBio 12 (2021).

69. E. Wong et al., The Vibrio cholerae colonization factor GbpA possesses a modular structure that governs binding to different host surfaces. PLoS Pathog 8, e1002373 (2012).

70. J. S. Loose, Z. Forsberg, M. W. Fraaije, V. G. Eijsink, G. Vaaje-Kolstad, A rapid quantitative activity assay shows that the Vibrio cholerae colonization factor GbpA is an active lytic polysaccharide monooxygenase. FEBS Lett 588, 3435–3440 (2014).

71. K. A. Syed et al., The Vibrio cholerae flagellar regulatory hierarchy controls expression of virulence factors. J Bacteriol 191, 6555–6570 (2009).

72. W. Song, J. A. Veenstra, N. Perrimon, Control of Lipid Metabolism by Tachykinin in Drosophila. Cell Rep 30, 2461 (2020).

73. A. J. Haugo, P. I. Watnick, *Vibrio cholerae* CytR is a repressor of biofilm development. Mol Microbiol 45, 471–483 (2002).

74. A. Dobin et al., STAR: ultrafast universal RNA-seq aligner. Bioinformatics 29, 15–21 (2013).

75. R. Patro, G. Duggal, M. I. Love, R. A. Irizarry, C. Kingsford, Salmon provides fast and bias-aware quantification of transcript expression. Nat Methods 14, 417–419 (2017).

76. M. I. Love, W. Huber, S. Anders, Moderated estimation of fold change and dispersion for RNA-seq data with DESeq2. Genome Biol 15, 550 (2014).

77. M. Cornwell et al., VIPER: Visualization Pipeline for RNA-seq, a Snakemake workflow for efficient and complete RNA-seq analysis. BMC Bioinformatics 19, 135 (2018).

